# General Methods for Evolutionary Quantitative Genetic Inference From Generalised Mixed Models

**DOI:** 10.1101/026377

**Authors:** Pierre de Villemereuil, Holger Schielzeth, Shinichi Nakagawa, Michael Morrissey

## Abstract

Methods for inference and interpretation of evolutionary quantitative genetic parameters, and for prediction of the response to selection, are best developed for traits with normal distributions. Many traits of evolutionary interest, including many life history and behavioural traits, have inherently non-normal distributions. The generalised linear mixed model (GLMM) framework has become a widely used tool for estimating quantitative genetic parameters for non-normal traits. However, whereas GLMMs provide inference on a statistically-convenient latent scale, it is sometimes desirable to express quantitative genetic parameters on the scale upon which traits are expressed. The parameters of a fitted GLMMs, despite being on a latent scale, fully determine all quantities of potential interest on the scale on which traits are expressed. We provide expressions for deriving each of such quantities, including population means, phenotypic (co)variances, variance components including additive genetic (co)variances, and parameters such as heritability. We demonstrate that fixed effects have a strong impact on those parameters and show how to deal for this effect by averaging or integrating over fixed effects. The expressions require integration of quantities determined by the link function, over distributions of latent values. In general cases, the required integrals must be solved numerically, but efficient methods are available and we provide an implementation in an R package, QGGLMM. We show that known formulae for quantities such as heritability of traits with Binomial and Poisson distributions are special cases of our expressions. Additionally, we show how fitted GLMM can be incorporated into existing methods for predicting evolutionary trajectories. We demonstrate the accuracy of the resulting method for evolutionary prediction by simulation, and apply our approach to data from a wild pedigreed vertebrate population.

## Introduction

Additive genetic variances and covariances of phenotypic traits determine the response to selection, and so are key determinants of the processes of adaptation in response to natural selection and of genetic improvement in response to artificial selection (Fisher, 1918; Falconer, 1960; Lynch and Walsh, 1998; Walsh and Lynch, forthcoming). While the concept of additive genetic variance (Fisher, 1918; Falconer, 1960) is very general, being applicable to any type of character with any arbitrary distribution, including, for example, fitness (Fisher, 1930), techniques for estimating additive genetic variances and covariances are best developed for Gaussian traits (i.e., traits that follow a normal distribution; Henderson 1950; Lynch and Walsh 1998). Furthermore, quantitative genetic theory for predicting responses to selection are also best developed and established for Gaussian characters (Walsh and Lynch, forthcoming), but see Morrissey (2015). Consequently, although many characters of potential evolutionary interest are not Gaussian (e.g. survival or number of offspring), they are not well-handled by existing theory and methods. Comprehensive systems for estimating genetic parameters and predicting evolutionary trajectories of non-Gaussian traits will hence be very useful for quantitative genetic studies of adaptation.

For the analysis of Gaussian traits, linear mixed model-based (LMM) inferences of genetic parameters, using the ‘animal model’, have become common practice in animal and plant breeding (Thompson, 2008; Hill and Kirkpatrick, 2010), but also in evolutionary studies on wild populations (Kruuk, 2004; Wilson *et al.*, 2010). Recently, the use of generalised linear mixed models (GLMMs) to analyse non-Gaussian traits has been increasing (e.g. Milot *et al.*, 2011; Wilson *et al.*, 2011; Morrissey *et al.*, 2012; de Villemereuil *et al.*, 2013; Ayers *et al.*, 2013). Whereas LMM analysis directly estimates additive genetic parameters on the scale on which traits are expressed and selected, and upon which we may most naturally consider their evolution, this is not the case for GLMMs. In this paper, we offer a comprehensive description of the assumptions of GLMMs and their consequences in terms of quantitative genetics and a framework to infer quantitative genetic parameters from GLMMs output. This work applies and extends theory in Morrissey (2015), to handle the effects of (non-linear) relationships among the scale upon which inference is conducted in a GLMM and the scale of data, and to accommodate the error structures that arise in GLMM analysis. These results generalise existing expressions for specific models (threshold model and Poisson with a log-link, Dempster and Lerner, 1950; Robertson, 1950; Foulley and Im, 1993). We show that fixed effects in GLMMs raise special complications and we offer some efficient approaches for dealing with this issue.

While it will undoubtedly be desirable to develop a comprehensive method for making data-scale inferences of quantitative genetic parameters with GLMMs, such an endeavour will not yield a system for predicting evolution in response to natural or artificial selection, even if a particular empirical system is very well served by the assumptions of a GLMM. This is because systems for evolutionary prediction, specifically the Breeder's equation (Lush, 1937; Fisher, 1924) and the Lande equation (Lande, 1979; Lande and Arnold, 1983), assume that breeding values (and in most applications, phenotypes) are multivariate normal or make assumptions such as linearity of the parent-offspring regression, which are unlikely to hold for non-normal traits (Walsh and Lynch, forthcoming). Even if it is possible to estimate additive genetic variances of traits on the scale upon which traits are expressed, we will show that these quantities will not strictly be usable for evolutionary prediction. However, we will see that the scale on which estimation is performed in a GLMM does, by definition, satisfy the assumptions of the Breeder's and Lande equations. Thus, for the purpose of predicting evolution, it may be useful to be able to express selection of non-Gaussian traits on this scale. Such an approach will yield a system for evolutionary prediction of characters that have been modelled with a GLMM, requiring no more assumptions than those that are already made in applying the statistical model.

The main results in this paper are arranged in four sections. First, we describe the GLMM framework: its relationship to the more general (Gaussian) LMM and especially to the Gaussian animal model (Henderson, 1973; Kruuk, 2004; Wilson *et al.*, 2010), how GLMMs can be usefully viewed as covering three scales and how some special interpretational challenges arise and are currently dealt with. Second, we propose a system for making inferences of quantitative genetic parameters on the scale upon which traits are expressed for arbitrary GLMMs. We show how to estimate genotypic and additive genetic variances and covariances on this scale, accounting for fixed effects as necessary. We lay out the formal theory underlying the system, apply it to an empirical dataset. The relationships between existing analytical formulae and our general framework are also highlighted. Third, we illustrate the issues when inferring quantitative genetic parameters using a GLMM with an empirical example on Soay sheep *(Ovis aries*) and how our framework can help to overcome them. Fourth, we outline a system of evolutionary prediction for non-Gaussian traits that capitalises on the fact that the latent scale in a GLMM satisfies the assumptions of available equations for the prediction of evolution. We show in a simulation study that (i) evolutionary predictions using additive genetic variances on the observed data scale represent approximations, and can, in fact, give substantial errors, and (ii) making inferences via the latent scale provides unbiased predictions, insofar as a GLMM may provide a pragmatic model of variation in non-Gaussian traits. The framework introduced here (including both quantitative genetic parameters inference and evolutionary prediction) has been implemeWhile it will undoubtedly be desirable to develop a comprehensive method for making data60
scale inferences of quantitative genetic parametersnted in a package for the R software (R Core Team, 2015) available at https://github.com/devillemereuil/qgglmm.

## The generalised linear mixed model framework

### Linear mixed models for gaussian traits

For Gaussian traits, a linear mixed model allows various analyses of factors that contribute to the mean and variance of phenotype. In particular, a formulation of a linear mixed model called the ‘animal model’ provides a very general method for estimating additive genetic variances and covariances, given arbitrary pedigree data, and potentially accounting for a range of different types of confounding variables, such as environmental effects, measurement error or maternal effects. A general statement of an animal model analysis decomposing variation in a trait, **z**, into additive genetic and other components would be

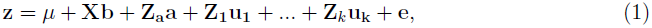

where *μ* is the model intercept, **b** is a vector of fixed effects such as sex and age, relating potentially both continuous and categorical effects to observations via the fixed effects design matrix **X**, just as in an ordinary linear model, and e is the vector of normally-distributed residuals. An arbitrary number of random effects can be modelled, with design matrices **Z**, where effects (**a, u_1_**…**u_k_**) are assumed to be drawn from normal distributions with variances to be estimated. The key feature of the animal model is that it includes individual additive genetic effects, or breeding values, conventionally denoted **a**. These additive genetic effects and, critically, their variance, are estimable given relatedness data, which can be derived from pedigree data, or, more recently, from genomic estimates of relatedness (Sillanpää, 2011). The covariances of breeding values among individuals can be modelled according to

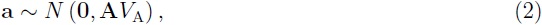

where **A** is the additive genetic relatedness matrix derived from the pedigree and *V*_A_ is the genetic additive variance.

### Common issues with non-gaussian traits

Many non-Gaussian traits, however, cannot be strictly additive on the scale on which they are expressed. Consider, for example, survival probability that is bounded at 0 and 1 so that effects like the substitution effect of one allele for another necessarily must be smaller when expressed in individuals that otherwise have expected values near zero or one. In such a scenario, it may be reasonable to assume that there exists an underlying scale, related to survival probability, upon which genetic and other effects are additive.

In addition to inherent non-additivity, many non-Gaussian traits will have complex patterns of variation. Over and above sources of variation that can be modelled with fixed and random effects, as in a LMM (e.g., using Eqs. 1 and 2), residual variation may include both inherently stochastic components, and components that correspond to un-modelled systematic differences among observations. In a LMM, such differences are not distinguished, but contribute to residual variance. However, for many non-Gaussian traits it may be desirable to treat the former as arising from some known statistical distribution, such as the binomial or Poisson distribution, and to deal with additional variation via a latent-scale residual (i.e. an overdispersion term). Separation of these two kinds of variation in residuals may be very generally useful in evolutionary quantitative genetic studies.

### The scales of the generalised linear mixed model

Generalised linear mixed model (GLMM) analysis can be used for inference of quantitative genetic parameters, and provides pragmatic ways of dealing with inherent non-additivity and with complex sources of variation. The GLMM framework can be thought of as consisting of three distinct scales on which we can think of variation in a trait occurring (see Fig. 1). A *latent scale* is assumed (Fig. 1, top), on which effects on the propensity for expression of some trait are assumed to be additive. A function, called a ‘link function’ is applied that links expected values for a trait to the latent scale. For example, a trait that is expressed in counts, say, number of behaviours expressed in a unit time, is a strictly non-negative quantity. As depicted in Fig. 1, a strictly positive distribution of expected values may related to latent values ranging from —∞ to +∞ by a function such as the log link. Finally, a distribution function (e.g. Binomial, Poisson, etc.) is required to model the “noise” of observed values around expected value (Fig. 1, bottom). Different distributions are suitable for different traits. For example, with a count trait such as that depicted in Fig. 1, observed values may be modelled using the Poisson distribution, with expectations related to the latent scale via the log link function.

More formally, these three scales of the GLMM can be written:

**Figure 1.**
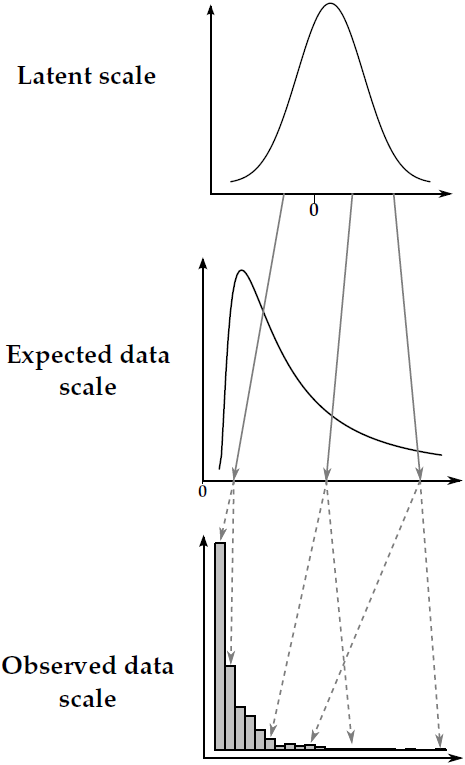
Example of the relationships between the three scales of the GLMM using a Poisson distribution and a logarithm link function. Deterministic relationships are denoted using grey plain arrows, whereas stochastic relationships are denoted using grey dashed arrows. Note that the latent scale is depicted as a simple Gaussian distribution for the sake of simplicity, whereas it is a mixture of Gaussian distributions in reality.

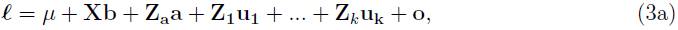

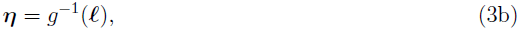

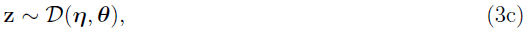

where Eq. 3a is just as for a LMM (Eq. 1), except that it describes variation on the *latent scale **l***, rather than the response directly. Note that we now refer to the “residual” (noted *e* in Eq. 1) as “overdispersion” (denoted **O**     , with a variance denoted *V*_O_), since residuals (variation around expected values) are defined by the distribution function, *D*, in this model. Just as for the LMM (Eq. 1), all random effects are assumed to follow normal distributions with variances to be estimated on the latent scale. Particularly, the variance of additive genetic effects a is assumed to follow Eq. 2 on the latent scale.

Eq. 3b formalises the idea of the link function. Any link function has an associated inverselink function, *g*^−1^, which is often useful for converting specific latent values to expected values. The expected values **η** constitute what we call the *expected data scale.* For example, where the log link function translates expected values to the latent scale, its inverse, the exponential function, translates latent values to expected values. Finally, Eq. 3c specifies the distribution by which the observations **z** scatter around the expected values according to some distribution function, that may involve parameters (denoted **θ**) other than the expectation. We call this the *observed data scale.* Some quantities of interest, such as the mean, are the same on the expected data scale and on the observed data scale. When parameters are equivalent on these two scales, we will refer to them together as the *data scales*.

The distinction we make between the expected and observed data scales is one of convenience as they are not different scales *per se.* However, this distinction allows for more biological subtlety when interpreting the output of a GLMM. The expected data scale can be thought of as the “intrinsic” value of individuals (shaped by both the genetic and the environment), but this intrinsic value can only be studied through random realisations. As we will see, because breeding values are intrinsic individual values, the additive genetic variance is the same for both scales, but, due to the added noise in observed data, the heritabilities are not. Upon which scale to calculate heritability depends on the underlying biological question. For example, individuals (given their juvenile growth and genetic value) might have an intrinsic annual reproductive success of 3.4, but can only produce a integer number of offspring each year (say 2, 3, 4 or 5): heritabilities of both intrinsic expectations and observed numbers can be computed, but their values and interpretations will differ.

### Current practices and issues to compute genetic quantitative parameters from GLMM outputs

Genetic variance components estimated in a generalised animal model are obtained on the latent scale. Hence, the “conventional” formula to compute heritability:

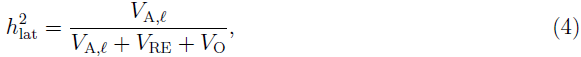

where *V*_RE_ is the summed variance of all random effects apart from the additive genetic variance, and *V*_O_ is the overdispersion variance, is the heritability on the latent scale, not on the observed data scale (Morrissey *et al.*, 2014). Here, and throughout this paper, *V*_A*l*_ stands for the additive genetic variance on the latent scale. Although it might sometimes be sensible to measure the heritability of a trait on the latent scale (for example, in animal breeding, where selection might be based on latent breeding values), it is natural to seek inferences on the scale upon which the trait is expressed, and on which we may think of selection as acting. Some expressions exist by which various parameters can be obtained or approximated on the observed data scale. For example, various expressions for the intra-class correlation coefficients on the data scale exist (reviewed in Nakagawa and Schielzeth, 2010), but, contrary to LMM, heritabilities on the data scales within a GLMM framework cannot be considered as intra-class correlation coefficients. Exact analytical expressions exist for the additive genetic variance and heritability on the observed data scale for two specific and important families of GLMMs (i.e. combinations of link functions and distribution functions): for a binomial model with a probit link function (i.e., the “threshold model,” Dempster and Lerner, 1950) and for a Poisson model with a logarithm link function (Foulley and Im, 1993). A general system for calculating genetic parameters on the expected and observed data scales for arbitrary GLMMs is currently lacking.

In addition to handling the relationship between observed data and the latent trait via the link and distribution functions, any system for expected and observed scale quantitative genetic inference with GLMMs will have to account for complex ways in which fixed effects can influence quantitative genetic parameters. It is currently appreciated that fixed effects in LMMs explain variance, and that variance associated with fixed effects can have a large influence on summary statistics such as repeatability (Nakagawa and Schielzeth, 2010) and heritability (Wilson, 2008). This principle holds for GLMMs as well, but fixed effects cause additional, important complications for interpreting GLMMs. While random and fixed effects are independent in a GLMM on the latent scale, the non-linearity of the link function renders them inter-related on the expected and observed scales. Consequently, and unlike in a LMM or in a GLMM on the latent scale, variance components on the observed scale in a GLMM depend on the fixed effects. Consider, for example, a GLMM with a log link function. Because the exponential is a convex function, the influence of fixed and random effects will create more variance on the expected and observed data scales for larger values than for smaller values.

## Quantitative genetic parameters in GLMMs

Although all examples and most equations in this article are presented in a univariate form, all our results are applicable to multivariate analysis, which is implemented in our software. Unless stated otherwise, the equations below assume that no fixed effect (apart from the intercept) were included in the GLMM model.

### Phenotypic mean and variances

**Expected population mean** The expected mean phenotype 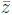 on the data scale (i.e., applying to both the mean expected value and mean observed value) is given by

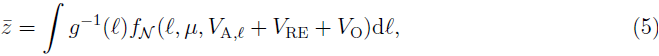

where *fN*(*l,μ*, *V*_A,*l*_,+*V*_RE_+*V*_o_) is the probability density of a Normal distribution with mean
μ and variance *V*_A,*l*_,+*V*_RE_+*V*_o_ evaluated at *l*.

**Expected-scale phenotypic variance** Phenotypic variance on the expected data scale can be obtained analogously to the data scale population mean. Having obtained 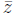, the phenotypic variance is

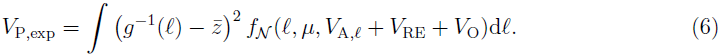

**Observed-scale phenotypic variance** Phenotypic variance of observed values is the sum of the variance in expected values and variance arising from the distribution function. Since these variances are independent by construction in a GLMM, they can be summed. This distribution variance is influenced by the latent trait value, but might also depend on additional distribution parameters included in **θ** (see Eq. 3c). Given a distribution-specific variance function *V*:

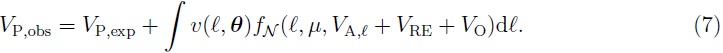

### Genotypic variance on the data scales, arising from additive genetic variance on the latent scale

Because the link function is non-linear, additive genetic variance on the latent scale is manifested as a combination of additive and non-additive variance on the data scales. Following Falconer (1960) the total genotypic variance on the data scale is the variance of genotypic values on that scale. Genotypic values are the expected data scale phenotypes, given latent scale genetic values. The expected phenotype of an individual with a given latent genetic value *a*, i.e., its genotypic value on the data scales *E*[*z*|*a*], is given by

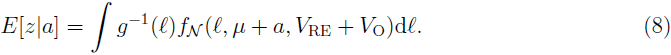

The total genotypic variances on the expected and observed data scales are the same, since genotypic values are expectations that do not change between the expected and observed scales. The total genotypic variance on both the expected and observed data scales is then

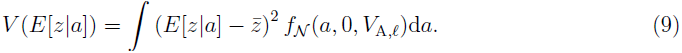

This is the total genotypic variance on the data scale, arising from strictly additive genetic variance on the latent scale. If non-additive genetic effects are modelled on the latent scale, they would be included in the expectations and integrals in Eqs. 8 and 9.

### Additive genetic variance on the data scales

The additive variance on the data scales is the variance of breeding values computed on the data scales. Following Robertson (1950; see also Fisher 1918), breeding values on the data scales, i.e., *a*_exp_ and *a*_obs_, are the part of the phenotype *z* that depends linearly on the latent breeding values. The breeding values on the datas scale can then be defined as the predictions of a least-squares regression of the observed data on the latent breeding values,

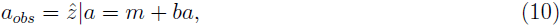

where 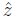 is the value of *z* predicted by the regression, a the latent breeding value and *m* and *b* the parameters of the regression. Thus, we have *V*_A,obs_ = *b*^2^*V*_A,*l*_ and, from standard regression theory:

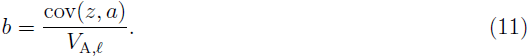

Because of the independence between the expected values of *z* (i.e. the expected data scale *g*^−*1*^ (*l*)) and the distribution “noise” (see Eq. 7), we can obtain the result that cov(*z,a*) = cov(g ^−1^(*l*),a), hence:

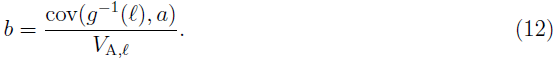

Stein's (1973) lemma states that if *X* and *Y* are bivariate normally distributed random variables, then the covariance of *Y* and some function of *X*, *f*(*X*), is equal to the expected value of *f* ′(X) times the covariance between *X* and *Y*, so,

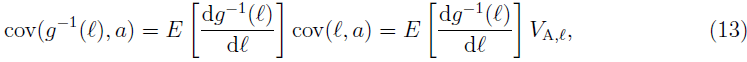

noting that the covariance of latent breeding values and latent values is the variance of latent breeding values. Finally, combining Eq. 12 with Eq. 13, we obtain:

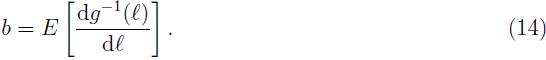

To avoid confusion with various uses of *b* as other forms of regression coefficients, and for consistency with Morrissey (2015), we denote the average derivative of expected value with respect to latent value as ψ:

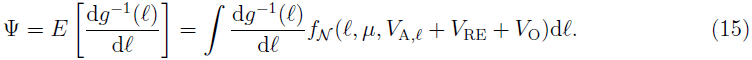

The additive genetic variance on the expected and observed scales are still the same and are given by

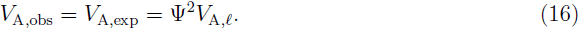

### Including fixed effects in the inference

**General issues** Because of the non-linearity introduced by the link function in a GLMM, all quantitative genetic parameters are directly influenced by the presence of fixed effects. Hence, when fixed effects are included in the model, it will often be important to marginalise over them to compute accurate population parameters. There are different approaches to do so. We will first describe the simplest approach (i.e. directly based on GLMM assumptions).

**Averaging over predicted values** In a GLMM, no assumption is made about the distribution of covariates in the fixed effects. Given this, we can marginalise over fixed effects by averaging over predicted values (marginalised over the random effects, i.e. 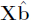, where 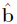 are the fixed effects estimates). Note that, doing so, we implicitly make the assumption that our sample is representative of the population of interest. Using this approach, we can compute the population mean in Eq. 5 as:

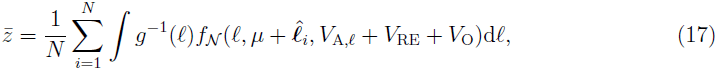

where *N* is the number of predicted latent values in 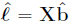. Typically, X will be the fixed effects design matrix used when fitting the generalised animal model (Eqs. 1, 2, and 3), and *N* will be the number of data observations. Furthermore, this assumes that all fixed effects represent biologically relevant variation, rather than being corrections for the observation process or experimental condition. From this estimate of 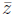, we can compute the expected-scale phenotypic variance:

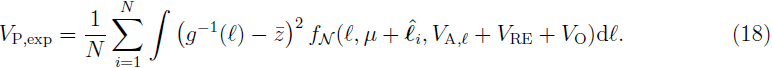

Note that we are not averaging over variances computed for each predicted values, since the value of the mean 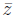 is the same across the computation. Eqs. 7, 8, 9 and 15 are to be modified accordingly to compute all parameters, including ψ. This approach has the advantages of being simple and making a direct use of the GLMM inference without further assumptions.

**Sampled covariates are not always representative of the population** The distribution of covariate values in **X** may not be representative of the population being studied. In such cases, integration over available values of fixed effects may be inappropriate. For example, a population may be known (or assumed) to have an equal sex ratio, but one sex may be easier to catch, and therefore over-represented in any given dataset. In such a situation, incorporation of additional assumptions or data about the distribution of covariates (e.g., of sex ratio) may be useful. A first approach is to predict values according to a new set of covariates constructed to be representative of the population (e.g. with balanced sex ratio). Given these new predicted values, the above approach can readily be used to compute quantitative genetic parameters of interest. A drawback of this approach is that it requires one to create a finite sample of predicted values instead of a full distribution of the covariates. A second approach will require one to specify such a distribution for fixed covariates, here noted *f*_*X*_(**X**). In that case, Eq. 17 can be modified as follows

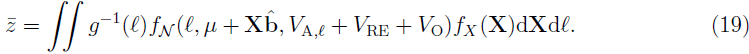

All relevant equations (Eqs. 6, 7, 8, 9 and 15) are to be modified accordingly. This approach is the most general one, but requires the ability to compute *f*_*X*_(**X**). Note that this distribution should also account for potential covariance between covariates.

### Summary statistics and multivariate extensions

Eqs. 5 through 16 give the values of different parameters that are useful for deriving other evolutionary quantitative genetic parameters on the observed data scale. Hence, from them, other parameters can be computed. The narrow-sense heritability on the observed data scale can be written as

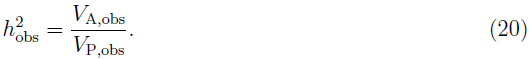

Replacing *V*_P,obs_ by *V*_P,exp_ will lead to the heritability on the expected data scale 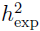

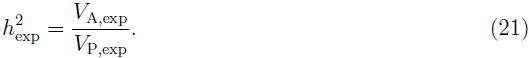

Recalling that *V*_A,obs_ = *V*_A,exp_, but *V*_P,obs_ ≠ *V*_P,exp_, note that the two heritabilities above differ. Parameters such as additive genetic coefficient of variance and evolvability (Houle, 1992) can be just as easily derived. The coefficient of variation on the expected and observed data scales are identical and can be computed as

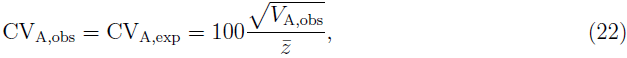

and the evolvability on the expected and observed data scales will be

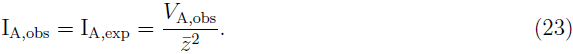

The multivariate genetic basis of phenotypes, especially as summarised by the **G** matrix, is also often of interest. For simplicity, all expressions considered to this point have been presented in univariate form. However, every expression has a fairly simple multivariate extension. Multivariate phenotypes are typically analysed by multi-response GLMMs. For example, the vector of mean phenotypes in a multivariate analysis on the expected data scale is obtained by defining the link function to be a vector-valued function, returning a vector of expected values from a vector of values on the latent scale. The phenotypic variance is then obtained by integrating the vector-valued link function times the multivariate normal distribution total variance on the latent scale, as in Eq. 5 and Eq. 7. Integration over fixed effects for calculation of the multivariate mean is directly analogous to either of the extensions of Eq. 5 given in Eqs. 17 or 19. Calculation of other parameters, such as multivariate genotypic values, additive-derived covariance matrices, and phenotypic covariance matrices, have directly equivalent multivariate versions as well. The additive genetic variance-covariance matrix (the **G** matrix) on the observed scale is simply the multivariate extension of Eq. 16, i.e., **G**_obs_ = **ψG**_*l*_**ψ**^*T*^. Here, **G**_*l*_ is the latent **G** matrix and **ψ** is the average gradient matrix of the vector-valued link function, which is a diagonal matrix of ψ values for each trait (simultaneously computed from a multivariate version of Eq. 15).

## Relationships with existing analytical formulae

### Binomial distribution and the threshold model

Heritabilities of binary traits have a long history of analysis with a threshold model (Wright, 1934; Dempster and Lerner, 1950; Robertson, 1950), whereby an alternate trait category is expressed when a trait on a latent “liability scale” exceeds a threshold. Note that this liability scale is not the same as the latent scale hereby defined for GLMM (see Fig. S1 in Supplementary Information). However, it can be shown (see Supplementary Information, section A) that a GLMM with a binomial distribution and a probit link function is exactly equivalent to such a model, only with slightly differently defined scales. For threshold models, heritability can be computed on this liability scale by using adding a so-called “link variance” *V*_L_ to the denominator (see for example Nakagawa and Schielzeth, 2010; de Villemereuil *et al.*, 2013):

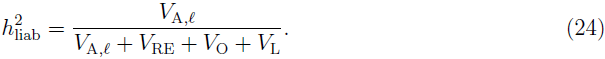

Because the probit link function is the inverse of the cumulative standard normal distribution function, the “link variance” *V*_L_ is equal to one in this case. One can think of the “link variance” as arising in this computation because of the reduction from three scales (in case of a GLMM) to two scales (liability and observed data in the case of a threshold model): the liability scale includes the link function.

When the heritability is computed using the threshold model, Dempster and Lerner (1950) and Robertson (1950) derived an exact analytical formula relating this estimate to the observed data scale:
_t_2

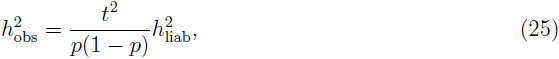

where *p* is the probability of occurrence of the minor phenotype and *t* is the density of a standard normal distribution at the *p*th quantile (see also Roff, 1997). It can be shown (see SI, section A) that this formula is an exact analytical solution to Eqs. 5 to 21 in the case of a GLMM with binomial distribution and a probit link. When fixed effects are included in the model, it is still possible to use these formulae by integration over the marginalised predictions (see SI, section A). Note that Eq. 25 applies only to analyses conducted with a probit link, it does not apply to a binomial model with a logit link function.

### Poisson distribution with a logarithm link

For a log link function and a Poisson distribution, both the derivative of the inverse link function, and the variance of the distribution, are equal to the expected value. Consequently, analytical results are obtainable for a log/Poisson model for quantities such as broad- and narrow-sense heritabilities. Foulley and Im (1993) derived an analytical formula to compute narrow-sense heritability on the observed scale:

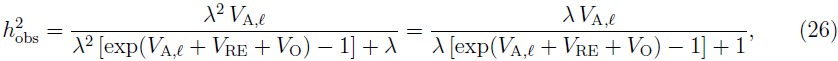

where λ is the data scale phenotypic mean, computed analytically as:

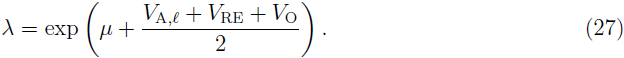

Again, it can be shown (see SI, section B) that these formulae are exact solutions to Eq. 5 to 21 when assuming a Poisson distribution with a log link. The inclusion of fixed effects in the model make the expression slightly more complicated (see SI, section B). These results can also be extended to the Negative-Binomial distribution with log link with slight modifications of the analytical expressions (see SI, section B).

The component of the broad-sense heritability on the observed data scale that arises from additive genetic effects on the latent scale can be computed as an intra-class correlation coefficient (i.e. repeatability) for this kind of model (Foulley and Im, 1993; Nakagawa and Schielzeth, 2010):

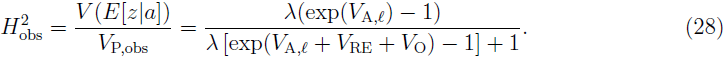

If non-additive genetic component were fitted in the model (e.g. dominance variance), they should be added to *V*_A,*l*_ in Eq. 28 to constitute the total genotypic variance, and thus obtain the actual broad-sense heritability. Note that the Eqs. 28 and 26 converge together for small values of *V*_A,*l*_.

## Example analysis: quantitative genetic parameters of a non-normal character

We modelled the first year survival of Soay sheep *(Ovis aries*) lambs on St Kilda, Outer Hebrides, Scotland. The data are comprised of 3814 individuals born between 1985 and 2011, and that are known to either have died in their first year, defined operationally as having died before the first of April in the year following their birth, or were known to have survived beyond their first year. Months of mortality for sheep of all ages are generally known from direct observation, and day of mortality is typically known. Furthermore, every lamb included in this analysis had a known sex and twin status (whether or not it had a twin), and a mother of a known age.

Pedigree information is available for the St Kilda Soay sheep study population. Maternal links are known from direct observation, with occasional inconsistencies corrected with genetic data. Paternal links are known from molecular data. Most paternity assignments are made with very high confidence, using a panel of 384 SNP markers, each with high minor allele frequencies, and spread evenly throughout the genome. Details of marker data and pedigree reconstruction are given in Bérénos *et al.* (2014). The pedigree information was pruned to include only phenotyped individuals and their ancestors. The pedigree used in our analyses thus included 4687 individuals with 4165 maternal links and 4054 paternal links.

We fitted a generalised linear mixed model of survival, with a *logit* link function and a binomial distribution function. We included fixed effects of individual's sex and twin status, and linear, quadratic, and cubic effects of maternal age (*matAge_i_*). Maternal age was mean-centred by subtracting the overall mean. We also included an interaction of sex and twin status, and an interaction of twin status with maternal age. We included random effects of breeding value (as for Eq. 2), maternal identity, and birth year. Because the overdispersion variance *V*_O_ in a binomial GLMM is unobservable for binary data, we set its variance to one. The model was fitted in MCMCglmm (Hadfield, 2010), with diffuse independent normal priors on all fixed effects, and parameter-expanded priors for the variances of all estimated random effects.

We identified important effects on individual survival probability, i.e., several fixed effects were substantial, and also, each of the additive genetic, maternal, and among-year random effects explained appreciable variance (Table 1). The model intercept corresponds to the expected value on the latent scale of a female singleton (i.e. not a twin) lamb with an average age (4.8 years) mother. Males have lower survival than females, and twins have lower survival than singletons. There were also substantial effects of maternal age, corresponding to a rapid increase in lamb survival with maternal age among relatively young mothers, and a negative curvature, such that the maximum survival probabilities occur among offspring of mothers aged 6 or 7 years. The trajectory of maternal age effects in the cubic model are similar to those obtained when maternal age is fitted as a multi-level effect.

**Table 1:** Parameters from the GLMM-based quantitative genetic analysis of Soay sheep *(Ovis aries*) lamb first-year survival. All estimates are reported as posterior modes with 95% credible intervals. The intercept in this model is arbitrarily defined for female lambs without twins, born to average age (4.8 years) mothers.

| Parameter | Posterior mode with 95% CI |
| --- | --- |
| (a) Fixed effects |  |
| Intercept | 2.686 (1.631 – 3.299) |
| Sex (male vs. female) | -1.185 (-1.436 – -0.932) |
| Twin (twin vs. singleton) | -2.383 (-3.315 – -1.760) |
| Maternal age, linear term | 0.238 (0.092 – 0.384) |
| Maternal age, quadratic term | -0.169 (-0.196 – -0.148) |
| Maternal age, cubic term | 0.014 (0.010 – 0.019) |
| Sex-twin interaction | 0.497 (0.016 – 1.016) |
| Sex-maternal age interaction | -0.020 (-0.103 – 0.070) |
| (b) Random effects |  |
| $V_{A,\ell}$ | 0.882 (0.256 – 1.542) |
| $V_{\text{mother}}$ | 0.467 (0.213 – 0.876) |
| $V_{\text{year}}$ | 3.062 (1.814 – 5.635) |

To illustrate the consequences of accounting for different fixed effects on expected and observed data scale inferences, we calculated several parameters under a series of different treatments of the latent scale parameters of the GLMM. We calculated the phenotypic mean, the additive genetic variance, the total variance of expected values, the total variance of observed values, and the heritability of survival on the expected and observed scales.

First, we calculated parameters using only the model intercept (μ in Eq. 1 and 3a). This intercept, under default settings, is arbitrarily defined by the linear modelling software implementation and is thus software-dependent. In the current case, due to the details of how the data were coded, the intercept is the latent scale prediction for female singletons with average aged (4.8 years) mothers. In an average year, singleton females with average aged mothers have a probability of survival of about 80%. The additive genetic variance *V*_A,obs_, calculated with Eq. 16 is about 0.005, and corresponds to heritabilities on the expected and observed scales of 0.115 and 0.042 (Table 2).

**Table 2:** Estimates of expected and observed scale phenotypic mean and variances, and additive genetic variance, for three different treatments of the fixed effects, as modelled on the linear scale with a GLMM, and reported in table 1. Additive genetic variance and heritability on the latent scales are also reported for comparison. Note that h2_at_ is slightly lower when averaging over fixed effects, since the variance they explain is then accounted for.

| Quantity | Arbitrary intercept<br>(singleton female) | Arbitrary intercept<br>(twin male) | Averaging over all fixed effects |
| --- | --- | --- | --- |
| $V_{A,\ell}$ | 0.915 (0.275 – 1.66) | 0.915 (0.275 – 1.66) | 0.915 (0.275 – 1.66) |
| $h_{\text{lat}}^2$ | 0.159 (0.056 – 0.27) | 0.159 (0.056 – 0.27) | 0.117 (0.0417 – 0.194) |
| $\bar{z}$ | 0.838 (0.721 – 0.887) | 0.352 (0.220 – 0.474) | 0.446 (0.334 – 0.511) |
| $V_{A, \text{obs}}$ | 0.006 (0.002 – 0.015) | 0.011 (0.005 – 0.023) | 0.013 (0.005 – 0.021) |
| $V_{P, \text{exp}}$ | 0.060 (0.034 – 0.095) | 0.098 (0.072 – 0.124) | 0.123 (0.106 – 0.137) |
| $V_{P, \text{obs}}$ | 0.136 (0.108 – 0.206) | 0.250 (0.183 – 0.250) | 0.250 (0.227 – 0.250) |
| $h_{\text{exp}}^2$ | 0.109 (0.043 – 0.201) | 0.122 (0.054 – 0.227) | 0.102 (0.039 – 0.166) |
| $h_{\text{obs}}^2$ | 0.047 (0.017 – 0.085) | 0.051 (0.022 – 0.101) | 0.043 (0.020 – 0.087) |

In contrast, if we wanted to calculate parameters using a different (but equally arbitrary) intercept, corresponding to twin males, we would obtain a mean survival rate of 0.32, an additive genetic variance that is twice as large, but similar heritabilities (Table 1). Note that we have not modelled any systematic differences in genetic parameters between females and males, or between singletons and twins. These differences in parameter estimates arise from the exact same estimated variance components on the latent scale, as a result of different fixed effects.

This first comparison has illustrated a major way in which the fixed effects in a GLMM influence inferences on the expected and observed data scales. For linear mixed models, it has been noted that variance in the response is explained by the fixed predictors, and that this may inappropriately reduce the phenotypic variance and inflate heritability estimates for some purposes (Wilson, 2008). However, in the example so far, we have simply considered two different intercepts (i.e. no difference in explained variance): female singletons vs male twins, in both cases, assuming focal groups of individuals are all born to average aged mothers. Again these differences in phenotypic variances and heritabilities arise from differences in intercepts, not any differences in variance explained by fixed effects. All parameters on the expected and observed value scales are dependent on the intercept, including the mean, the additive genetic variance and the total variance generated from random effects. Heritability is modestly affected by the intercept, because additive genetic and total variances are similarly, but not identically, influenced by the model intercept.

Additive genetic effects are those arising from the average effect of alleles on phenotype, integrated over all background genetic and environmental circumstances in which alternate alleles might occur. Fixed effects, where they represent biologically-relevant variation, are part of this background. Following our framework (see Eq. 17), we can solve the issue of the influence of the intercept by integrating our calculation of ψ and ultimately *V*_A,obs_ over all fixed effects. This approach has the advantage of being consistent for any chosen intercept, as the value obtained after integration will not depend on that intercept. Considering all fixed and random effects, quantitative genetic parameters on the expected and observed scales are given in table 2, third column. Note that additive genetic variance is not intermediate between the two extremes (concerning sex and twin status), that we previously considered. The calculation of *V*_A,obs_ now includes an average slope calculated over a wide range of the steep part of the inverse-link function (near 0 on the latent scale, and near 0.5 on the expected data scale), and so is relatively high. The observed total phenotypic variance Vp_;obs_ is also quite high. The increase in *V*_P,obs_ has two causes. First the survival mean is closer to 0.5, so the random effects variance is now manifested as greater total variance on the expected and observed scales. Second, there is now variance arising from fixed effects that is included in the total variance.

Given this, which estimates should be reported or interpreted? We have seen that when fixed effects are included in a GLMM, the quantitative genetic parameters calculated without integration are sensitive to an arbitrary parameter: the intercept. Hence integration over fixed effects may often be the best strategy for obtaining parameters that are not arbitrary. In the case of survival analysed here, 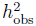 is the heritability of realised survival, whereas 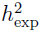 is the heritability of “intrinsic” individual survival. Since realised survival is the one “visible” by natural selection, 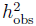 might be a more relevant evolutionary parameter. Nonetheless, we recommend that *V*_P,exp_ and *V*_P,obs_ are both reported.

## Evolutionary prediction

Systems for predicting adaptive evolution in response to phenotypic selection assume that the distribution of breeding values is multivariate normal, and in most applications, that the joint distribution of phenotypes and breeding values is multivariate normal (Lande, 1979; Lande and Arnold, 1983; Morrissey, 2014; Walsh and Lynch, forthcoming). The distribution of breeding values is assumed to be normal on the latent scale in a GLMM analysis, and therefore the parent-offspring regression will also be normal on that scale, but not necessarily on the data scale. Consequently, evolutionary change predicted directly using data-scale parameters may be distorted. The Breeder's and Lande equations may hold approximately, and may perhaps be useful. However, having taken up the non-trivial task of pursuing GLMM-based quantitative genetic analysis, the investigator has at their disposal inferences on the latent scale. On this scale, the assumptions required to predict the evolution of quantitative traits hold. In this section we will first demonstrate by simulation how application of the Breeder's equation will generate biased predictions of evolution. We then proceed to an exposition of some statistical machinery that can be used to predict evolution on the latent scale (from which evolution on the expected and observed scale can subsequently be calculated, using Eq. 5), given inference of the function relating traits to fitness.

### Direct Application of the Breeder's and Lande equations on the data scale

In order to explore the predictions of the Breeder's equation applied at the level of observed phenotype, we conducted a simulation in which phenotypes were generated according to a Poisson GLMM (Eq. 3a to 3c, with a Poisson distribution function and a log link function), and then selected the largest observed count values (positive selection) with a range of proportions of selected individuals (from 5% to 95%, creating a range of selection differentials), a range of latent-scale heritabilities (0.1, 0.3, 0.5 and 0.8, with a latent phenotypic variance fixed to 0.1), and a range of latent means *μ* (from 0 to 3). We simulated 10,000 replicates of each scenario, each composed of a different array of 10,000 individuals. For each simulation, we simulated 10,000 offspring. For each offspring, a breeding value was simulated according to *a_*l*,*i*_* ∼ *N((a_l,d_*=*a_l,s_)/,V_A,*l*_/2)*, where *a_l,i_* is the focal offspring's breeding value, *a_*l*,d_* and *a*_l,s_ are the breeding values of simulated dams and sires and *V*_A,*l*_/2 represents the segregational variance assuming parents are not inbred. Dams and sires were chosen at random with replacement from among the pool of simulated selected individuals. For each scenario, we calculated the realised selection differential arising from the simulated truncation selection, *S*_obs_, and the average evolutionary response across simulations, *R*_obs_. For each scenario, we calculated the heritability on the observed scale using Eq. 20. If the Breeder's equation was strictly valid for a Poisson GLMM on the observed scale, the realised heritability *R*_obs_/*S*_obs_ would be equal to the observed-scale heritability 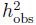.

The correspondence between *R*_obs_/*S*_obs_ and 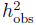 is approximate (Fig. 2), and strongly depends on the selection differential (controlled here by the proportion of selected individuals). Note that, although the results presented here depict a situation where the ratio R_obs_/S_obs_ is very often larger than 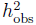, this is not a general result (e.g. this is not the case when using negative instead of positive selection, data not shown). In particular, evolutionary predictions are poorest in absolute terms for large *μ* and large (latent) heritabilities. However, because we were analysing simulation data, we could track the selection differential of latent value (by calculating the difference in its mean between simulated survivors and the mean simulated before selection). We can also calculate the mean latent breeding value after selection. Across all simulation scenarios, the ratio of the change in breeding value after selection, to the change in breeding value before selection was equal to the latent heritability (see Fig. 2), showing that evolutionary changes could be accurately predicted on the latent scale.

**Figure 2.**
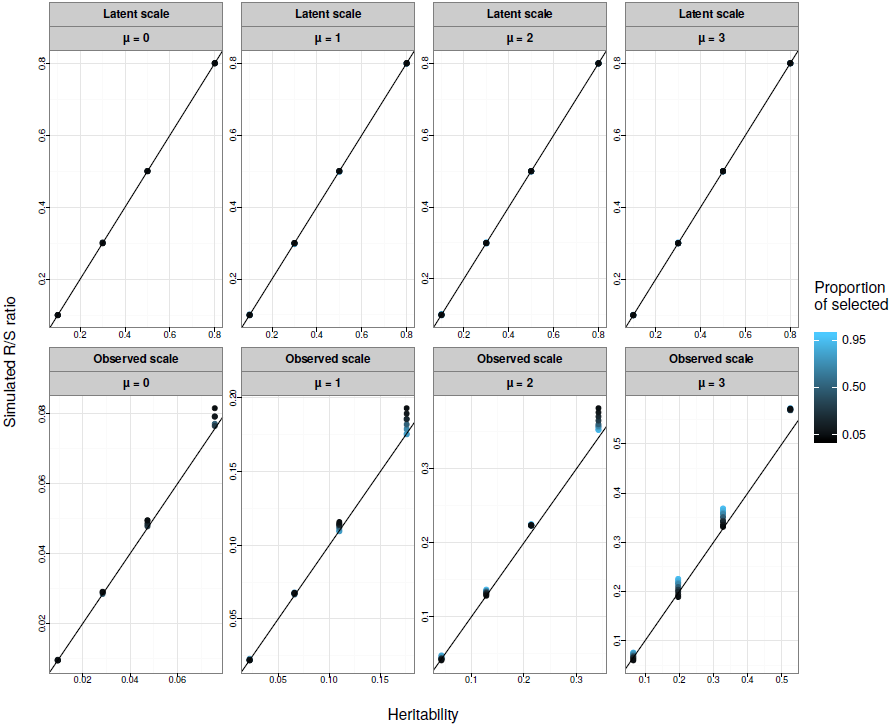
Simulated R/S (evolutionary response over selection differential, or the realised heritability) on the latent (upper panels) or observed data (lower panels) scales against the corresponding-scale heritabilities. Each data point is the average over 10,000 replicates of 10,000 individuals for various latent heritabilities 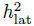 (0.1, 0.3, 0.5, 0.8), latent population mean (μ from 0 to 3, from left to right) and proportion of selected individuals (5%, 10%, 20%, 30%, 50%, 70%, 80%, 90%, 95%, varying from black to blue). The 1:1 line is plotted in black. The breeder's equation is predictive on the latent scale (upper panels), but approximate on the observed data scale (lower panels), because phenotypes and breeding values are not jointly multivariate normal on that scale.

### Evolutionary change on the latent scale, and associated change on the expected and observed scales

In an analysis of real data, latent (breeding) values are, of course, not measured. However, given an estimate of the effect of traits on fitness, say via regression analysis, we can derive the parameters necessary to predict evolution on the latent scale. The idea is thus to relate measured fitness on the observed data scale to the latent scale, compute the evolutionary response on the latent scale and finally compute the evolutionary response on the observed data scale.

To relate the measured fitness on the observed scale to the latent scale, we need to compute the expected fitness *W*_exp_ given latent trait value *l*, which is

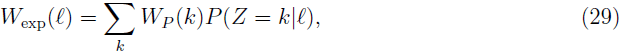

where *W_P_(k)* is the measure of fitness for the *k*th data scale category (assuming the observed data scale is discrete as in most GLMMs). Population mean fitness, can then be calculated in an analogous way to Eq. 5:

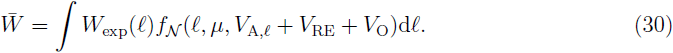

These expressions comprise the basic functions necessary to predict evolution. Given a fitted GLMM, and a given estimate of the fitness function *W_P_(k)*, each of several approaches could give equivalent results. For simplicity, we proceed via application of the breeder's equation at the level of the latent scale.

The change in the mean genetic value of any character due to selection is equal to the covariance of breeding value with relative fitness (Robertson, 1966, 1968). Using Stein's (1973) lemma once more, this covariance can be obtained as the product of the additive genetic variance of latent values and the average derivative of expected fitness with respect to latent value, i.e., 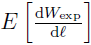.Evolution on the latent scale can therefore be predicted by

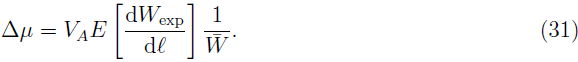

In the case of a multivariate analysis, note that the derivative above should be a vector of partial derivatives (partial first order derivative with respect to latent value for each trait) of fitness.

If fixed effects need to be considered, the approach can be modified in the same way as integration over fixed effects is accomplished for calculating other quantities, i.e. the expression

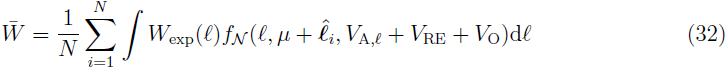

would be used in calculations of mean fitness and the average derivative of expected fitness with respect to latent value.

Phenotypic change caused by changes in allele frequencies in response to selection is calculated as

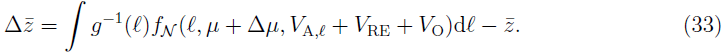

Or, if fixed effects are included in the model:

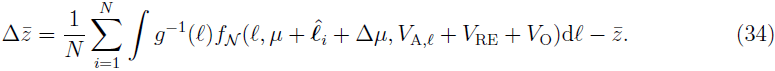

Note that, in this second equation, 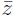 must be computed as in Eq. 17 and that this equation assumes that the distribution of fixed effects for the offspring generation is the same as for the parental one.

Another derivation of the expected evolutionary response using the Price-Robertson identity (Robertson, 1966; Price, 1970) is given in the Supplementary Information (section C).

### The simulation study revisited

Using the same replicates as in the simulation study above, we used Eqs. 29 to 34 to predict phenotypic evolution. This procedure provides greatly improved predictions of evolutionary change on the observed scale (Fig. 3, top row). However, somewhat less response to selection is observed than is predicted. This deviation occurs because, in addition to producing a permanent evolutionary response in the mean value on the latent scale, directional selection creates a transient reduction of additive genetic variance due to linkage disequilibrium. Because the link function is non-linear, this transient change in the variance on the latent scale generates a transient change in the mean on the expected and observed scales. Following several generations of random mating, the evolutionary change on the observed scale would converge on the predicted values. We simulated such a generation at equilibrium by simply drawing breeding values for the post-selection sample from a distribution with the same variance as in the parental generation. This procedure necessarily generated a strong match between predicted and simulated evolution (Fig. 3, bottom row). Additionally, the effects of transient reduction in genetic variance on the latent scale could be directly modelled, for example, using Bulmer's (1971) approximations for the transient dynamics of the genetic variance in response to selection.

**Figure 3.**
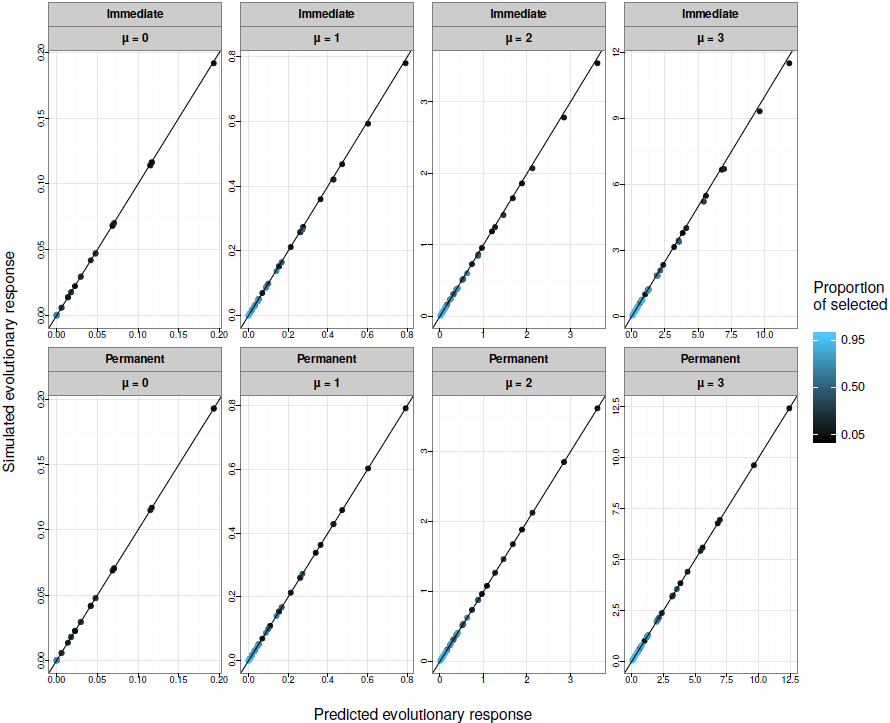
Predicted *R*_obs_ (phenotypic evolutionary response on the observed scale, see Eq. 34) against the simulated *R*_obs_, via evolutionary predictions applied on the latent scale. Each data point is the average over 10,000 replicates of 10,000 individuals for various latent heritabilities 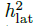 (0.1, 0.3, 0.5, 0.8), latent population mean (μ from 0 to 3) and proportion of selected individuals (5%, 10%, 20%, 30%, 50%, 70%, 80%, 90%, 95%, varying from black to blue). The 1:1 line is plotted in black. The upper panels (“Immediate”) show simulations for the response after a single generation, which include both a permanent and transient response to selection arising from linkage disequilibrium. The bottom panels (“permanent”) show simulation results modified to reflect only the permanent response to selection.

## Discussion

The general approach outlined here for quantitative genetic inference with GLMMs has several desirable features: *(i)* it is a general framework, which should work with any given GLMM and especially, any link and distribution function, *(ii)* provides mechanisms for rigorously handling fixed effects, which can be especially important in GLMMs, and *(iii)* it can be used for evolutionary prediction under standard GLMM assumptions about the genetic architecture of traits.

Currently, with the increasing applicability of GLMMs, investigators seem eager to convert to the observed data scale. It seems clear that conversions between scales are generally useful. However, it is of note that the underlying assumption when using GLMMs for evolutionary prediction is that predictions hold on the latent scale. Hence, some properties of heritabilities for additive Gaussian traits, particularly the manner in which they can be used to predict evolution, do not hold on the data scale for non-Gaussian traits, even when expressed on the data scale. Yet, given an estimate of a fitness function, no further assumptions are necessary to predict evolution on the data scale, via the latent scale (as with Eqs. 29, 31, and 33), over and above those that are made in the first place upon deciding to pursue GLMM-based quantitative genetic analysis. Hence we recommend using this framework to produce accurate predictions about evolutionary scenarios.

We have highlighted important ways in which fixed effects influence quantitative genetic inferences with GLMMs, and developed an approach for handling these complexities. In LMMs, the main consideration pertaining to fixed effects is that they explain variance, and some or all of this variance might be inappropriate to exclude from an assessment of *V_P_* when calculating heritabilities (Wilson, 2008). This aspect of fixed effects is relevant to GLMMs, but furthermore, all parameters on the expected and observed scales, not just means, are influenced by fixed effects in GLMMs; this includes additive genetic and phenotypic variances. This fact necessitates particular care in interpreting GLMMs. Our work clearly demonstrates that consideration of fixed effects is essential, and the exact course of action needs to be considered on a case-by-case basis. Integrating over fixed effects would solve, in particular, the issue of intercept arbitrariness illustrated with the Soay sheep example. Yet cases may often arise where fixed effects are fitted, but where one would not want to integrate over them (e.g. because they represent experimental rather than natural variability). In such cases, it will be important to work with a biologically meaningful intercept, which can be achieved for example by centring covariates on relevant values (Schielzeth, 2010). Finally, note that this is not an all-or-none alternative: in some situations, it could be relevant to integrate over some fixed effects (e.g. of biological importance) while some other fixed effects (e.g. those of experimental origins) would be left aside.

One of the most difficult concepts in GLMMs seen as a non-linear developmental model (Morrissey, 2015) is that an irreducible noise is attached to the observed data. This is the reason why we believe that distinguishing between expected and observed data scale does have a biological meaning. Researchers using GLMMs need to realise that this kind of model can assume a large variance in the observed data with very little variance on the latent and expected data scales. For example, a Poisson/log GLMM with a latent mean *μ* = 0 and a total latent variance of 0.5 will result in observed data with a variance *V*_P,obs_ = 2.3. Less than half of this variance lies in the expected data scale (*V*_P,exp_ = 1.07), the rest is residual Poisson variation. Our model hence assumes that more than half of the measured variance comes from totally random noise. Hence, even assuming that the whole latent variance is composed of additive genetic variance, the heritability will never reach a value above 0.5. Whether this random noise should be accounted for when computing heritabilities (i.e. whether we should compute 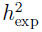 or 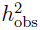) depends on the evolutionary question under study. In many instances, it is likely that natural selection will act directly on realised values rather than their expectations, in which case 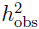 should be preferred. We recommend however, that, along with *V*_A,obs_, all other variances (including *V_o_*, *V*_P,exp_ and *V*_P,obs_) are reported by researchers.

The expressions given here for quantitative genetic parameters on the expected and observed data scales are exact, given the GLMM model assumptions, in two senses. First, they are not approximations, such as might be obtained by linear approximations (Browne *et al.*, 2005). Second, they are expressions for the parameters of direct interest, rather than convenient substitutes. For example, the calculation (also suggested by Browne *et al.* 2005) of variance partition coefficients (i.e. intraclass correlations) on an underlying scale only provides a value of the broad-sense heritability (e.g. using the genotypic variance arising from additive genetic effects on the latent scale).

The framework developed here (including univariate and multivariate parameters computation and evolutionary predictions on the observed data scale) is implemented in the R package **QGGLMM** available at https://github.com/devillemereuil/qgglmm. Note that the package does not perform any GLMM inference but only implements the hereby introduced framework for analysis posterior to a GLMM inference. While the calculations we provide will often (i.e. when no analytical formula exists) be more computationally demanding than calculations on the latent scale, they will be direct ascertainments of specific parameters of interest, since the scale of evolutionary interest is likely to be the observed data scale, rather than the latent scale (unless some artificial selection is applied to predicted latent breeding values as in modern animal breeding). Most applications should not be onerous. Computations of means and (additive genetic) variances took less than a second on a 1.7 GHz processor when using our R functions on the Soay sheep data set. Summation over fixed effects, and integration over 1000 posterior samples of the fitted model took several minutes. When analytical expressions are available (e.g. for Poisson/log, Binomial/probit and Negative-Binomial/log, see the supplementary information and R package documentation), these computations are considerably accelerated.

## Acknowledgements

PdV was supported by a doctoral studentship from the French *Ministère de la Recherche et de l'Enseignement Superieur.* HS was supported by an Emmy Noether fellowship from the German Research Foundation (DFG; SCHI 1188/1-1). SN is supported by a Future Fellowship, Australia (FT130100268). MBM is supported by a University Research Fellowship from the Royal Society (London). The Soay sheep data were provided by Josephine Pemberton and Loeske Kruuk, and were collected primarily by Jill Pilkington and Andrew MacColl with the help of many volunteers. The collection of the Soay sheep data is supported by the National Trust for Scotland and QinetQ, with funding from NERC, the Royal Society, and the Leverhulme Trust. We thank Kerry Johnson, Paul Johnson, Alastair Wilson, Loeske Kruuk and Josephine Pemberton for valuable discussions and comments on this manuscript.

